# Defining new Buruli ulcer endemic areas in urban southeastern Australia using bacterial genomics-informed possum excreta surveys

**DOI:** 10.1101/2025.07.29.667575

**Authors:** Andrew H. Buultjens, Koen Vandelannoote, Jessica L. Porter, Stephen Muhi, Emma C. Hobbs, Catarina Agostinho Antão, EE Laine Tay, Jake Lacey, Norelle Sherry, Maria Globan, Caroline J. Lavender, Anna Meredith, Paul D. R. Johnson, Katherine B. Gibney, Timothy P. Stinear

## Abstract

Buruli ulcer (BU) in southeastern Australia is a zoonosis caused by infection with *Mycobacterium ulcerans*. Australian native possums are a major wildlife reservoir and infected possums shed *M. ulcerans* in their excreta, with mosquitoes being the major transmitting vector in this region. BU is geographically restricted and this feature, combined with an average 4.8-month incubation period, makes tracking *M. ulcerans* environmental spread and timely identification of new BU endemic areas challenging. While human movement complicates transmission tracing, we used the highly territorial behaviour of native possums and high-resolution pathogen genomics to confidently identify new BU endemic areas in Melbourne’s inner northwest and southern suburbs of Geelong. Using pathogen genomic phylodynamic modelling, we estimated that *M. ulcerans* was introduced to these areas 2-6 years before the emergence of human BU cases. This study shows how possum excreta surveys combined with pathogen genome data can pinpoint new BU endemic areas, thus providing critical local knowledge for targeted public health interventions to reduce exposure risk and ensure early diagnosis.

## INTRODUCTION

Buruli ulcer (BU) is a disfiguring disease caused by *Mycobacterium ulcerans*, a slow-growing bacterium that produces mycolactone, a potent immunosuppressive and cytotoxic toxin (1–4). First described in the 1930s in southeastern Australia (5), BU causes extensive tissue necrosis and long-term disability, though it is rarely fatal (6). While the transmission route remained unclear for decades, recent studies have confirmed that mosquitoes transmit *M. ulcerans* to humans in Australia (7), though transmission elsewhere remains poorly understood.

The epidemiology of BU is highly focal, with endemic and non-endemic areas often only kilometres apart (4). BU has been reported in over 30 countries (8), including regions in Asia, the Western Pacific, the Americas and across West and Central Africa (9). In Australia, most cases occur in the state of Victoria (in southeastern Australia), though BU has also been reported in the Northern Territory (9), Queensland (10–12) and New South Wales (13, 14). Historical Victorian hotspots include Gippsland (15), Phillip Island (16) and Frankston/Langwarrin (17), with recent endemicity on the Mornington and Bellarine Peninsulas (18–20) (Figure 1). Since becoming notifiable in 2004, detailed case data has been available (21), but the long and variable incubation period (mean 4.8 months, IQR 101–171 days) complicates exposure location inference, especially in multi-endemic regions (22, 23). While global cases are declining, BU incidence is rising in Victoria, with both sustained increases and geographic expansion in southeastern Australia (24–26).

**Figure 1.**
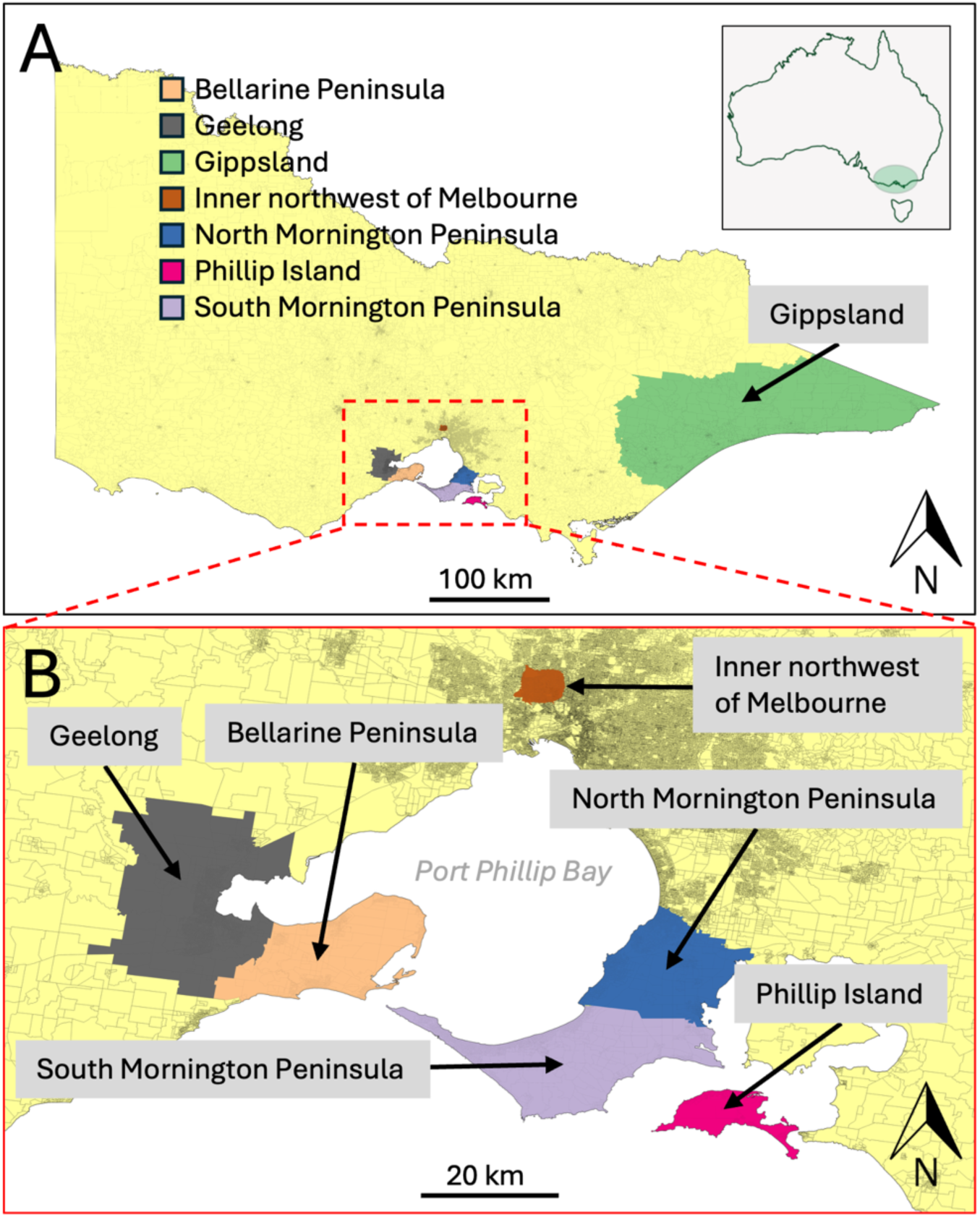
Map of major BU endemic areas in southeastern Australia. (A) Regional context of the BU endemic areas within the state of Victoria and inset shows national context; (B) BU endemic areas within the Port Phillip Bay region. The basemap depicts population density (boundaries of 2011 census data meshblocks).

A key challenge in studying *M. ulcerans* ecology is its extreme difficulty to culture from environmental samples, likely due to overgrowth by faster-growing microbes (27). However, sensitive qPCR surveys have detected high concentrations of M. ulcerans DNA in the faeces of common ringtail (*Pseudocheirus peregrinus*) and, to a lesser extent, common brushtail (*Trichosurus vulpecula*) possums, suggesting they serve as wildlife reservoirs (28). This association is so strong that models using possum excreta data have successfully predicted future human BU risk (29).

Over the past two decades, bacterial population genomics has advanced our understanding of *M. ulcerans* evolution, pathogenicity and epidemiology (7, 30–33). In southeastern Australia, genomic data suggest a westward spread from a Gippsland ancestor (33), with SNP signatures in clinical isolates unique to specific endemic areas (7, 33). These likely reflect population bottlenecks followed by clonal expansion and newly emerging endemic areas are therefore expected to show distinct SNP profiles. Recent culture-independent sequencing has recovered whole *M. ulcerans* genomes from *IS2404*-positive possum excreta and mosquito DNA, which matched clinical genomes, confirming a shared transmission pathway (7).

Given the highly localised nature of BU transmission and the rise in cases beyond well-established endemic zones, there is an urgent need to identify emerging areas in real time. Early detection enables timely public health interventions to disrupt transmission, raise community awareness and alert frontline clinicians to consider BU in diagnoses, ultimately improving patient outcomes (26). Detecting local transmission is challenging in urban settings due to high human mobility and complex exposure histories. In contrast, possums have small home ranges (∼100 m) (34), making their excreta a reliable spatial marker of pathogen presence. With genomic typing of possum-derived *M. ulcerans* genomes, clinical bacterial isolates can be compared to local strains, improving source attribution.

In this study, we investigated two newly emerged BU endemic areas in Melbourne’s inner northwest and Geelong’s southern suburbs by combining possum faecal surveys with microbial genomics, demonstrating the value of genomics-informed environmental surveillance for confirming local transmission.

## METHODS

### Study sites

The area of study in Melbourne’s inner northwest is located approximately eight kilometres northwest of Melbourne’s Central Business District (CBD) (Victoria’s capital city) in the state of Victoria, Australia. The area incorporates the suburbs of Essendon (population 21,240), Brunswick West (population 14,318), Pascoe Vale South (population 10,390), Moonee Ponds (population 16,224) and Strathmore (population 10,031) (35). The Geelong South area borders the Geelong CBD (Victoria’s second largest city, southwest of Melbourne) and is bound to the south by the Barwon River. The Geelong South study area is centred on the suburb of Belmont (population 42,257) (35). Both regions are predominantly residential suburban land use with some light industry. Native possums have adapted well to suburban gardens and live close to humans throughout Melbourne and Geelong.

### Possum excreta sampling

Field sampling in Melbourne’s inner northwest and Geelong was performed along 200 m intervals as previously described (29). The 2021 and 2022 surveys in Melbourne’s inner northwest and 2022 Geelong survey were conducted as part of this research while the Geelong 2020 survey was previously published (36).

### *IS2404* quantitative PCR

Quantitative PCR (qPCR) screening targeting the *IS2404* high copy insertion sequence was performed as described (29). Cycle threshold (Ct) values of 40 and above were classified as ‘not detected’ as they are considered false positives.

### GIS mapping

All GIS mapping was conducted with QGIS (v3.26.2-Buenos Aires). Shapefile census 2011 data was downloaded from the Australian Bureau of Statistics (www.abs.gov.au/AUSSTATS/abs@.nsf/DetailsPage/1270.0.55.001July%202011).

### Ethics

Ethical approval for the use of de-identified human BU case location data, aggregated at the mesh block level, was granted by the Victorian Department of Health Human Ethics Committee (HREC/54166/DHHS-2019-179235).

### BU case notification data

The Victorian Department of Health (DH) provided a de-identified dataset of all BU cases notified in Victoria during 2020-2022. A BU case was defined as a patient with a clinically suggestive lesion and laboratory confirmation of *M. ulcerans* DNA via *IS2404* real-time PCR (37, 38). BU has been a notifiable communicable disease in Victoria since 2004. Presumed exposure locations were reported at the mesh block level - the highest-resolution unit in Australian census data, typically encompassing 30 - 60 dwellings. Cases were included in this analysis if the reported exposure location fell within the inner northwest suburbs of Melbourne (bounding box: −37.747062°, 144.890000° to −37.762315°, 144.951622°) or the Geelong region (bounding box: −38.073318°, 144.274189° to −38.207718°, 144.418465°) and if symptom onset date fell within the incubation period interquartile range (101 - 171 days) of the possum excreta sampling period - that is, symptom onset occurred between 101 and 171 days after the start and end of sampling, respectively.

### Spatial clustering

Spatial clustering was performed using SatScan (39) as described previously for both *M. ulcerans IS2404* PCR-positivity in excreta samples (Bernoulli model) and using mesh blocks with zero to multiple human BU cases (Poisson model) (29). Files used to perform this analysis have been made available on a GitHub repository (https://github.com/abuultjens/BU-endemic-area-detection).

### Genome sequencing

Mycobacterial culture and whole genome sequencing was performed as described (33). Clinical isolate genomes from BU cases from 2017 through to 2023 with DH reported exposure in Melbourne’s inner northwest and the southern suburbs of Geelong were sequenced for this study. To provide phylogenomic context, we included 45 previously published *M. ulcerans* genomes isolated from 1945 to 2016 from key endemic regions across southeastern Australia, including Gippsland, Phillip Island, the Bellarine Peninsula and the Mornington Peninsula. Whole-genome sequencing of *IS2404* PCR-positive possum excreta specimens was conducted using a hybridisation capture method, as previously described, for samples collected in Melbourne’s inner northwest and Geelong’s southern suburbs (7). Sequence reads generated from this study were submitted to the National Centre for Biotechnology Information (NCBI) GenBank and listed in Table S1 (available in GitHub repository: https://github.com/abuultjens/BU-endemic-area-detection).

### Bioinformatic analysis

Illumina paired DNA sequence reads were mapped against a fully assembled *M. ulcerans* Victorian clinical isolate chromosome using Snippy (v4.4.5) (JKD8049; GenBank accession NZ_CP085200.1; https://github.com/tseemann/snippy). All clinical isolate genomes and the sequence enrichment datasets were processed using the default Snippy minimum coverage threshold of 10x. However, to accommodate the lower coverage of the sequence capture datasets, a reduced threshold of 1x was applied. To minimise the inclusion of false-positive SNPs, the core SNPs identified among the clinical isolate genomes were used to filter the sequence capture SNPs, retaining only those higher confidence variants (minimum of 10x coverage) in the final core SNP alignment. A phylogenomic tree was inferred by maximum likelihood using FastTree (v.2.1.10) with the General Time Reversible model of nucleotide substitution (40) from an alignment of core genome SNPs obtained by Snippy, as above.

### Phylodynamic molecular clock analyses

BEAST2 v2.7.7 was used to both infer a time-tree of Victorian *M. ulcerans* isolates and estimate key divergence times. BEAUti v2.7.7 input consisted of an alignment of all SNPs in the non-repetitive core genome. Tip-dates were defined as the time of strain isolation and entered with varying precision based on the availability of exact calendar dates. bModelTest v1.3.3 (41) was used to average over (marginalise) the site model in the phylogenetic analysis. The uncorrelated log-normal relaxed molecular clock and a coalescent Bayesian Skyline Plot (BSP) tree prior were selected (42). BEAUti xml files were supplemented with the number of invariant sites in the genome by manually specifying the amount of invariant A, C, T and G nucleotides. Analysis was performed in BEAST2 using four independent chains of 100 million Markov chain Monte Carlo (MCMC) generations, with samples taken every 10,000 MCMC generations. The xml file used in the BEAST2 analysis is available in file S1. Log files were checked in Tracer v.1.7.2 (43) for proper mixing and convergence and to assess whether the used chain length produced an effective sampling size (ESS) larger than 400 for all parameters (which indicates sufficient sampling). LogCombiner v2.7.7 was then used to combine tree and log files of the four independent BEAST2 runs, after removing a 10% burn-in from each run. TreeAnnotator v2.7.7 was used to summarise the posterior sample of time-trees into a maximum clade credibility tree. The reliability of our estimates of divergence times and timescales was assessed by conducting a date-randomisation test (44). This test consists of repeating the BEAST2 analysis 20 times with identical model settings with randomly reshuffled sampling dates. Date randomisation of the BEAST2 xml file was performed using the R package Tipdatingbeast v1.1-0 (45) in R v4.3.3.

## RESULTS

### Emergence of new BU endemic areas in Melbourne and Geelong

In 2017, BU cases began to emerge among residents of several inner northwestern suburbs of Melbourne, areas with no prior history of local *M. ulcerans* transmission (Figure 1). Similarly, in the southern suburbs of Geelong, there had been a notable increase in cases from 2019, with cases first reported from the area in 2011 (36) (Figure 1). Initially, these infections were presumed to have been acquired during travel to nearby historical and well-documented BU endemic areas in the region. However, epidemiological analysis that included assessment of BU case travel histories and comparison of residential addresses revealed geographic clustering, suggesting these cases from suburban areas of Melbourne’s inner northwest and Geelong represented local transmission (Figure 1).

### Detection of *M. ulcerans* DNA in possum excreta to confirm local transmission

We have previously shown that possum excreta testing using qPCR screening is a sensitive method for detecting the environmental presence of *M. ulcerans*, which is associated with local transmission to humans (29). Here, we found that *M. ulcerans* was reliably detectable in possum excreta collected with median *IS2404* qPCR Ct values of 36.59, 33.89 and 36.14 for the Melbourne’s inner northwest 2021, 2022 and Geelong 2022 surveys, respectively (Figure S1).

### *M. ulcerans* presence in possum excreta and disease in humans overlap in time and space

The spatial distribution of *M. ulcerans* in possum excreta and its correlation with human BU cases was assessed through two different field surveys in Melbourne’s inner northwest (2021 and 2022) and two surveys in Geelong (2020 and 2022). Scanning spatial analysis using the location of *IS2404* qPCR-positive and negative possum excreta from these surveys identified non-random clusters of *M. ulcerans* positive possum excreta (Figure 2). In all four surveys there was spatial overlap between clusters of *M. ulcerans* positive possum excreta and clusters of human BU cases. Positive possum excreta samples were consistently found near areas where human BU cases occurred. The overlapping clusters were present across both regions and each year from 2020 to 2022, inclusive. This pattern was most evident in Melbourne’s inner northwestern in the suburbs of Brunswick West, Pascoe Vale South and Essendon (Figure 2 A and B), as well as in Geelong around the suburb of Belmont (Figure 2 C, D and E).

**Figure 2.**
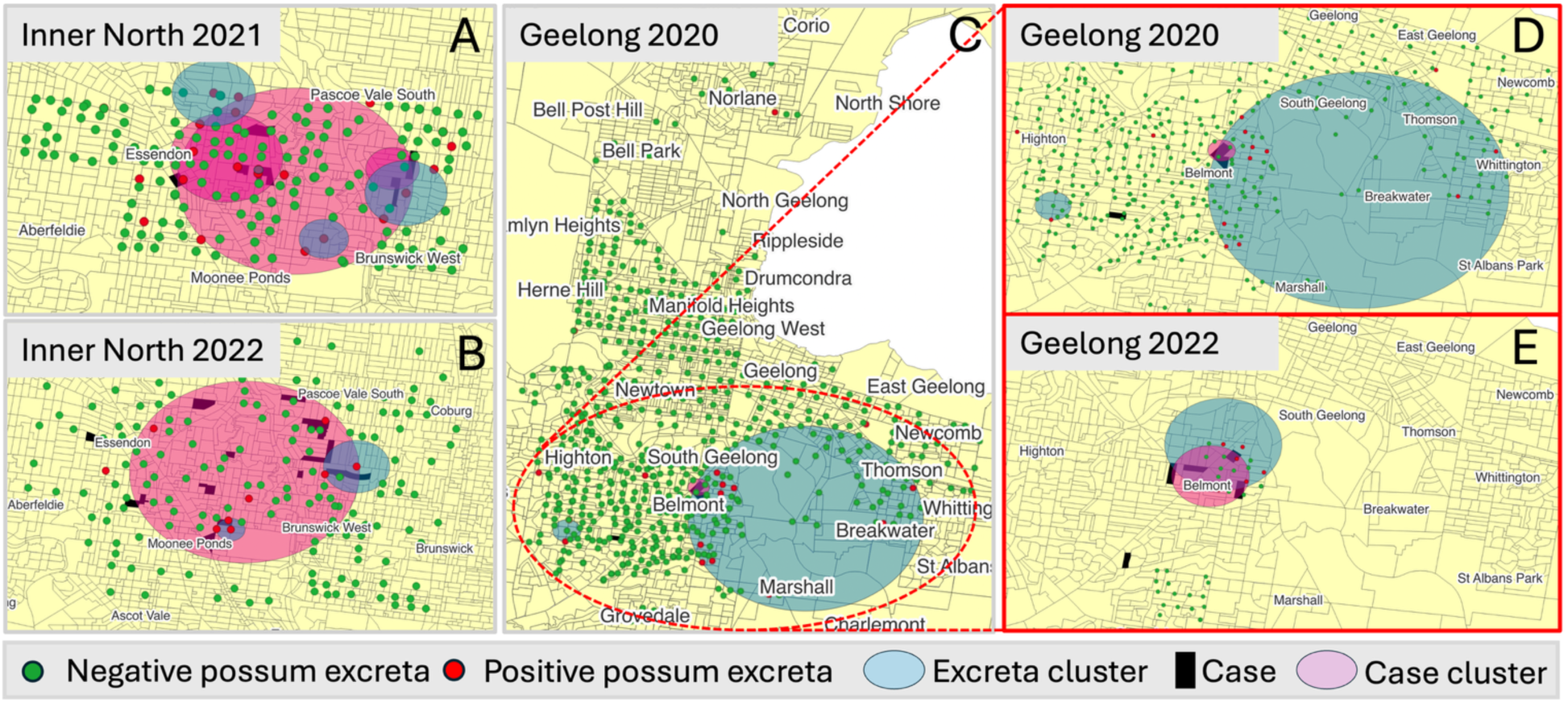
Spatial distribution of possum excreta samples and human BU cases. Possum excreta samples are shown (red are IS2404 qPCR-positive and green are negative), with SatScan clusters for M. ulcerans presence in possum excreta (light blue) and human BU cases (light red). Mesh blocks with human cases are shown in solid black. A: Melbourne’s inner northwest, 2021. B: Melbourne’s inner northwest, 2022. C: Geelong-wide survey, 2020. D: Zoomed-in view of Belmont area, Geelong, 2020. E: Zoomed-in view of Belmont area, Geelong, 2022.

### Phylodynamic modelling provides temporal estimates for the founding *M. ulcerans* **ancestors of newly emerged endemic areas**

Mapping of bacterial genome sequence reads against a local *M. ulcerans* reference chromosome revealed 218 evenly distributed core genome SNPs (Figure 3A). These genomes were selected to capture the spatial and temporal diversity of the pathogen across all major endemic regions of Victoria, including cases dating back to 1945 in Gippsland (5). A phylodynamic analysis was performed using the resulting core genome SNP alignment to investigate the population structure of clinical isolates from Melbourne’s inner northwest and the Geelong region. The phylogenetic tree (Figure 3B) includes 102 *M. ulcerans* genomes obtained from 43 clinical isolates representing the broader population diversity across southeastern Australia, 37 isolates from Melbourne’s inner northwest and 22 isolates from Geelong. *M. ulcerans* clinical isolate genomes from Melbourne’s inner northwest formed a well-supported monophyletic cluster within the Victorian phylogeny, consistent with localised transmission and a recent common origin. In contrast, isolate genomes from the Geelong region exhibited greater genomic diversity, forming two distinct sub-lineages (hereafter referred to as Geelong-1 and Geelong-2) within the broader Victorian phylogeny (Figure 3B).

**Figure 3.**
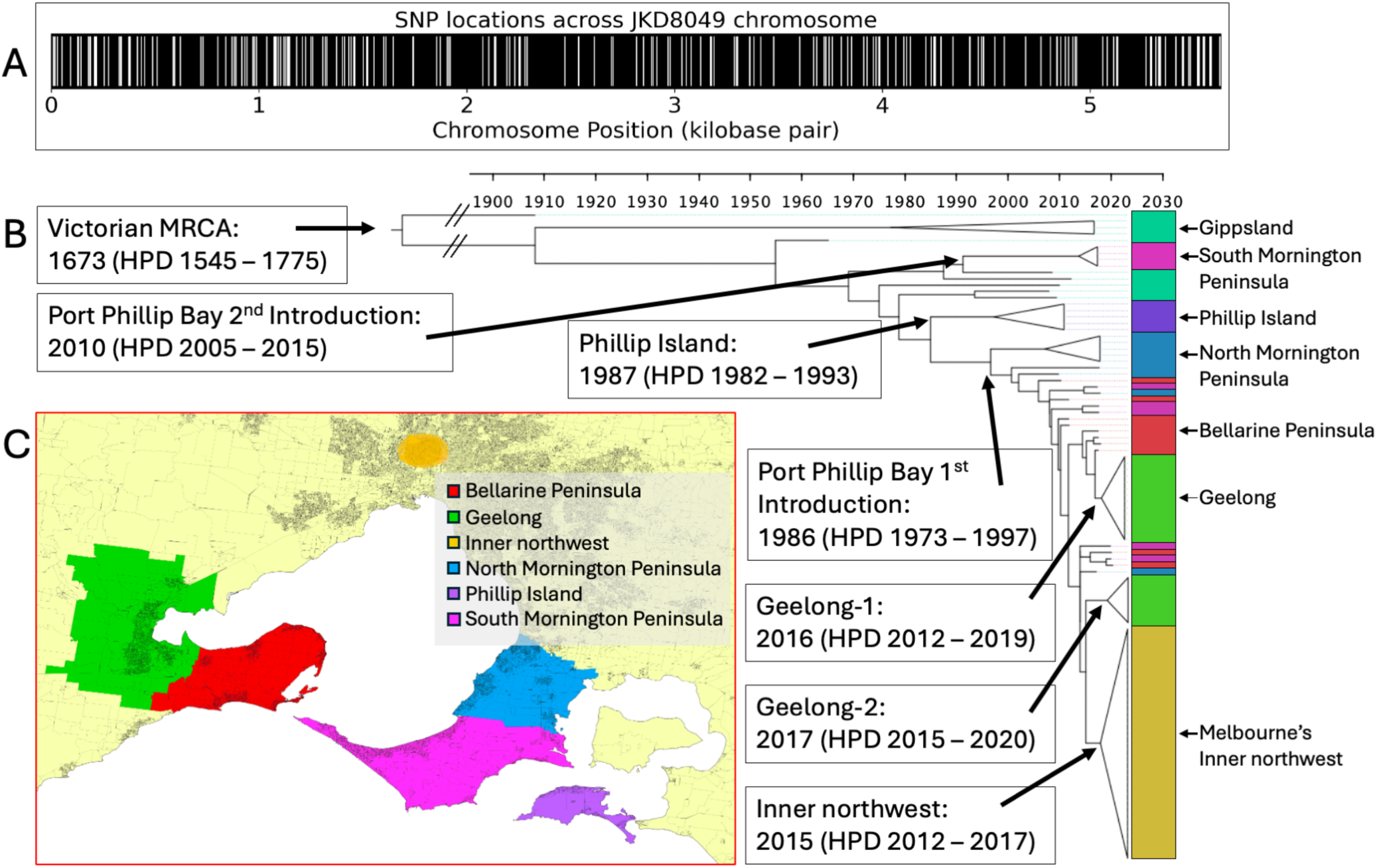
Phylodynamic analysis of M. ulcerans isolates from southeastern Australia. (A) Distribution of 218 core SNPs across the M. ulcerans reference chromosome, derived from mapping clinical isolate genome sequence reads to a local, finished reference genome. (B) Time-calibrated phylogenomic tree showing the population structure of M. ulcerans clinical isolates from Melbourne’s inner northwest and Geelong, placed within a broader context of temporally and geographically diverse Victorian clinical isolates (tree built using 218 core SNPs). Major lineages are annotated with estimated dates of the most recent common ancestors. Lineages that are geographically monophyletic (comprising isolates exclusively from a specific endemic region) have been collapsed into clade triangles. Branch lengths reflect time, as indicated by the scale. (C) Map of Victoria indicating BU-endemic areas corresponding to the isolates included in the phylogeny, with colours matched to isolate labels in panel B.

Using culture isolation dates to calibrate a molecular clock, we estimated the temporal emergence of key ancestral nodes in the phylogeny. The most recent common ancestor (MRCA) of Melbourne’s inner northwest cluster was dated to approximately 2015 (95% highest posterior density [HPD]: 2012-2017), with cases in that region first reported in 2017 (2 years after the ancestral temporal estimate). The Geelong-1 and Geelong-2 lineages were estimated to have emerged in 2016 (HPD: 2012-2019) and 2017 (HPD: 2015-2020), respectively, with cases in that region first reported in 2011 and a notable uptick in case from 2019. For comparison, the MRCA of the Phillip Island cluster was dated to 1987 (HPD: 1982-1993) with first cases reported in 1993 (16) (6 years later), while two distinct Port Phillip Bay MRCAs were estimated to have emerged in 1986 (HPD: 1973-1997) and 2010 (HPD: 2005-2015), with the first Port Phillip Bay cases reported in 1990 (17) (4 years after the ancestral temporal estimate of the first introduction). The overall MRCA for the *M. ulcerans* population in southeastern Australia was estimated to have emerged around 1673 (HPD: 1545-1775).

### Phylogenomic structure of *M. ulcerans* in Melbourne’s inner northwest and Geelong endemic areas

To specifically examine the genomic diversity among Melbourne’s inner northwest and Geelong lineages, we generated focused maximum likelihood trees using SNP alignments generated using only genomes obtained from *M. ulcerans* patient isolates from those individual locations. The phylogeny inferred using Melbourne’s inner northwest genomes revealed a population structure that aligns with specific exposure locations within this area. In particular, the Brunswick West and Pascoe Vale South genomes appeared to lie near the base of the sub-tree, suggesting they represent some of the earliest sampled forms of this lineage, with subsequent pathogen spread into the Essendon area, followed by further spread and genetic diversification into Moonee Ponds. (Figure 4 A and B). The only evidence of geographical structure among the Geelong isolates was a subtree defined by four SNPs within the Geelong-1 lineage, which included two clinical isolate genomes from Newtown (Figure 4 C and D).

**Figure 4.**
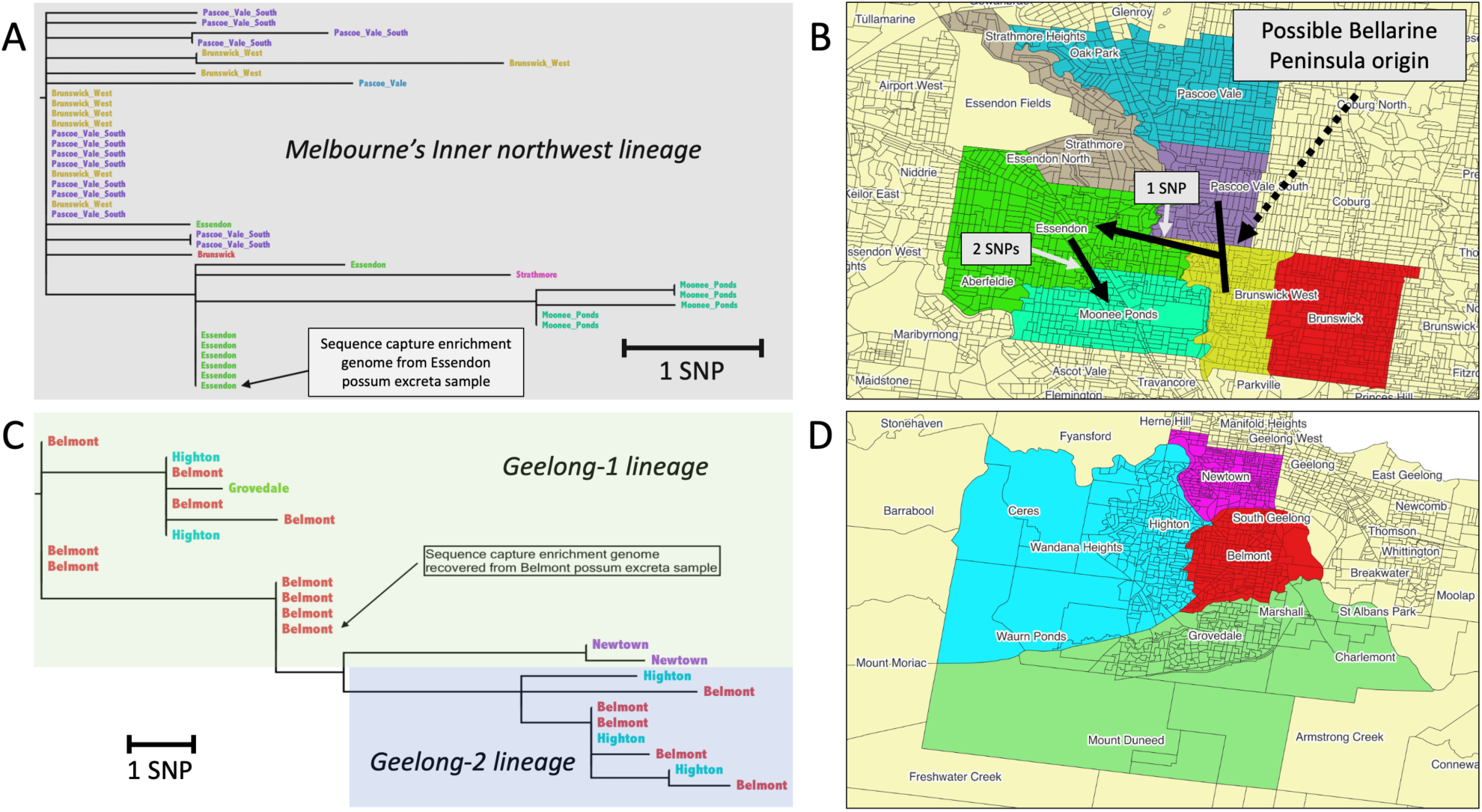
Phylogenomic and spatial analysis of M. ulcerans isolates from two newly identified BU-endemic areas in Victoria. (A, C) Maximum likelihood phylogenetic trees of M. ulcerans clinical isolate genomes from Melbourne’s inner northwest (22 core SNPs) and the Geelong region (18 core SNPs), respectively. M. ulcerans genomes recovered from possum excreta are indicated. Branch lengths reflect core genome SNP differences, as indicated by the scale. The genomes detected in possum excreta are indicated in both trees. (B, D) Maps of Melbourne’s inner northwest and Geelong, respectively, showing the suburbs of likely patient exposure for the isolate genomes included in panels A and C, respectively. Suburbs are colour-coded to match the corresponding tip labels in the phylogenetic trees.

Using sequence capture enrichment, we successfully recovered *M. ulcerans* whole genome sequence data from possum excreta in each of the newly identified endemic areas and incorporated these data into the core genome phylogeny alongside clinical isolate genomes. The *M. ulcerans* genome recovered from possum excreta from Essendon, in Melbourne’s inner northwest, clustered tightly within a cluster of Essendon clinical isolates, providing strong genomic evidence of a shared transmission pathway between possums and humans in that area (Figure 4A). Similarly, the *M. ulcerans* genome recovered from possum excreta from a Belmont sample grouped within the Geelong-1 subtree among Belmont clinical isolates, again consistent with local pathogen transmission (Figure 4C).

### Model of Melbourne’s inner northwest geographical spread of *M. ulcerans*

Based on the phylogenomic evidence, a model of the geographical spread of *M. ulcerans* in Melbourne’s inner northwest was proposed (Figure 4B). The pathogen appears to have been firstly introduced into the Brunswick West and Pascoe Vale South areas, possibly originating from the Bellarine Peninsula, a well-documented endemic region (26), as the basal inner northwest genotypes are indistinguishable from those originating in Barwon Heads (33). From there, *M. ulcerans* has likely spread westward into Essendon, with a subtree defined by one SNP (position 470,691 of JKD8049 reference chromosome). As the *M. ulcerans* population continued to spread from Essendon, it then likely moved further south into Moonee Ponds, where a further distinct cluster emerged, defined by an additional two SNPs (positions 3,981,241 and 5,294,886).

## DISCUSSION

Buruli ulcer (BU) transmission is highly localised, with established links between possums, mosquitoes and humans in southeastern Australia (7, 28, 29, 46). As BU cases rise and new endemic areas emerge, including Melbourne’s inner northwest and Geelong’s southern suburbs (26), we investigated these expansions using structured possum excreta surveys and pathogen genomic analyses.

Due to *M. ulcerans’* long incubation period (mean 4.8 months) and frequent exposure to multiple endemic areas, patient recall is an unreliable indicator of infection origin. To address this, we compared *M. ulcerans* genomes from possum excreta (a spatially trustworthy analyte) with those from BU patients. Previous studies showed spatial and temporal overlap between possum and human cases, with matching genomes confirming shared transmission (7, 29). Here, we extend these findings to define and confirm newly emerged BU endemic areas.

Unique SNP patterns indicate recent evolutionary bottlenecks following pathogen introduction and expansion. We analysed *M. ulcerans* genomes from BU patients and possum excreta in Melbourne’s inner northwest and Geelong, using phylodynamic methods to assess population structure and estimate emergence dates, revealing clear genotype-geography associations. *M. ulcerans* genomes from the two newly emerged endemic areas showed a distinctive clustering between possum-derived and clinical isolate genomes, consistent with local transmission and patterns seen in other endemic areas (7). Phylogeographic analysis suggests that *M. ulcerans* was introduced into Brunswick West and Pascoe Vale South around 2015, likely from the Bellarine Peninsula, followed by spread to Essendon and Moonee Ponds, where a distinct cluster emerged. *M. ulcerans* genomes from BU cases in Geelong formed two distinct clusters (Geelong-1 and Geelong-2), with a possum-derived genome falling within Geelong-1 and likely reflect two separate introductions around 2016 and 2017, both potentially originating from the Bellarine Peninsula. During this study, the discovery of an inner northwest-specific genomic cluster was reported to the Victorian Department of Health, prompting a targeted health advisory with prevention messages and guidance for local clinicians to consider BU in patients exposed to the area (47–49).

Phylodynamic modelling estimated *M. ulcerans* emergence in Melbourne’s inner northwest occurring 2 - 6 years (median 4.5) before BU cases, suggesting a lag due to expansion in possum reservoirs before spillover (33). Geelong is an exception, with lineage emergence dating after initial cases in 2011 - likely reflecting misattributed exposures or isolated introductions. In contrast, Geelong-1 and Geelong-2 appear to reflect later introductions that drove local transmission, consistent with increased cases from 2019 (36). Combined genomic and environmental sampling highlight the power of pathogen genomics to resolve introductions and spread within a small urban area (eight square kilometres in this study). Recent genomic analyses also identified local transmission in Batemans Bay, New South Wales, where two non-travelling BU cases had clinical isolates forming a distinct *M. ulcerans* clade, further demonstrating the utility of genomics in identifying emerging endemic areas (14).

The Geelong possum-derived *M. ulcerans* genome clustered within the Geelong-1 lineage, suggesting further sampling could uncover Geelong-2 genotypes and highlighting the need to increase sequencing from positive possum faecal specimens. However, sequence capture enrichment is expensive ($500 AUD/sample) and limited to high-DNA samples (Ct ≤ 25), while all but two positive specimens in this study had Ct > 25. To address this, we are developing a low-cost ($80 AUD) PCR-based amplicon sequencing method targeting informative SNPs (manuscript in preparation), which will enable broader genomic surveillance and clarify environmental transmission by comparing SNPs across hosts and vectors.

The mechanism of long-distance spatial spread of *M. ulcerans* remains unclear. However, recent findings that 30% of fox faecal samples in southeastern Australia were *IS2404*-positive with viable bacteria (50) and that foxes prey on possums (51, 52), suggest a possible role in pathogen dissemination that merits further investigation.

## CONCLUSION

The ongoing geographical expansion of BU endemicity highlights the need for both sensitive and specific methods to identify emerging areas to provide timely warnings to local communities and institute measures to reduce spread to humans. The recent discovery that mosquitoes vector *M. ulcerans* to humans now offers a clear prevention strategy by avoiding mosquito bites, underscoring the importance of early, targeted community messaging to disrupt transmission. Possum excreta surveys detect *M. ulcerans*, indicating potential endemicity, while genomic analyses provide specificity by linking human disease to the local environment through genome matching, collectively confirming local transmission (7, 47–49). This integrated approach - connecting wildlife, environmental surveillance and human health - is a strong example of a One Health strategy. Therefore, future investigations into the emergence of novel BU endemic areas would benefit from combining possum excreta surveys with pathogen genomics.

## ACKNOWLEDGEMENTS

We thank Joe Gleeson, Jeremy Bourke, Oscar Howden, and Chris Sanders for assistance with fieldwork.

