## Supplementary material for "Defining new Buruli ulcer endemic areas in urban southeastern Australia using bacterial genomics-informed possum excreta surveys": Figure S1

**Supplementary figures:**

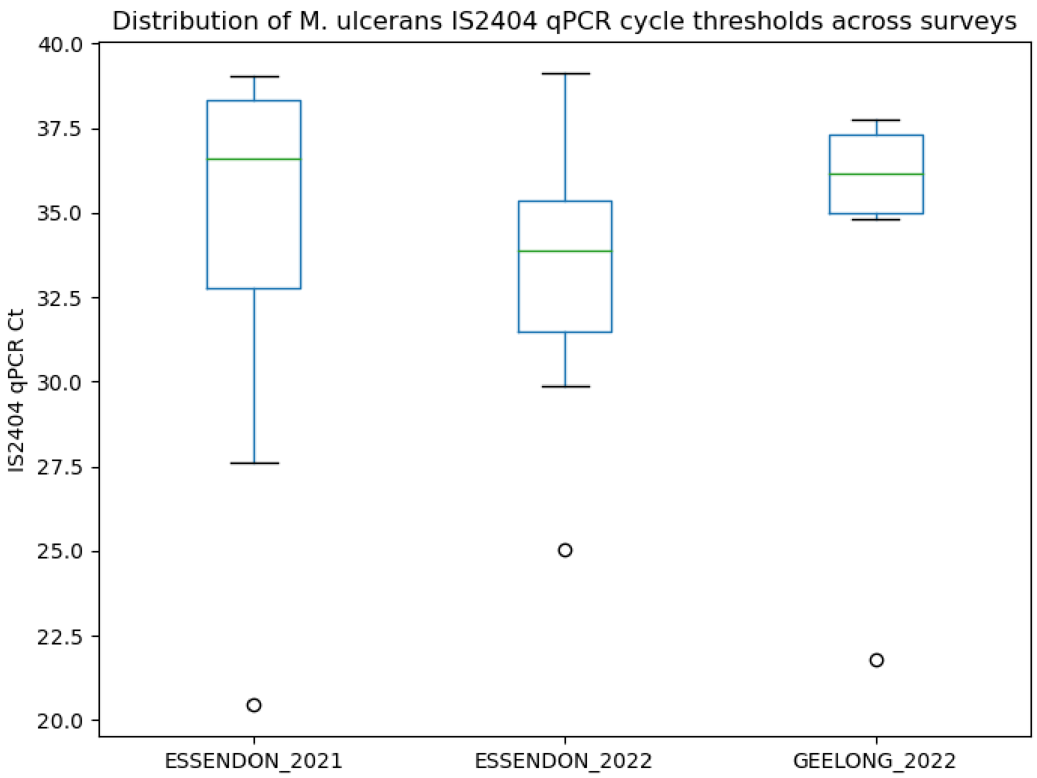


**Figure. S1** *Distribution of IS2404 qPCR cycle thresholds for possum excreta across different surveys and sites. Boxplot error bars represent the range within 1.5 x IQR of the lower and upper quartiles.*
